# Design of Specific Primer Set for Detection of B.1.1.7 SARS-CoV-2 Variant using Deep Learning

**DOI:** 10.1101/2020.12.29.424715

**Authors:** Alejandro Lopez-Rincon, Carmina A. Perez-Romero, Alberto Tonda, Lucero Mendoza-Maldonado, Eric Claassen, Johan Garssen, Aletta D. Kraneveld

**Affiliations:** Division of Pharmacology, Utrecht Institute for Pharmaceutical Sciences, Faculty of Science, Utrecht University, Universiteitsweg 99, 3584 CG Utrecht, the Netherlands; Departamento de Investigación, Universidad Central de Queretaro (UNICEQ), Av. 5 de Febrero 1602, San Pablo, 76130 Santiago de Querétaro, Qro., Mexico; UMR 518 MIA-Paris, INRAE, c/o 113 rue Nationale, 75103, Paris, France; Hospital Civil de Guadalajara “Dr. Juan I. Menchaca”. Salvador Quevedo y Zubieta 750, Independencia Oriente, C.P. 44340 Guadalajara, Jalisco, México; Department of Viroscience, Erasmus Medical Center, Rotterdam, the Netherlands; Athena Institute, Vrije Universiteit, De Boelelaan 1085, 1081 HV Amsterdam, the Netherlands; Department Immunology, Danone Nutricia research, Uppsalalaan 12, 3584 CT Utrecht, the Netherlands

## Abstract

The SARS-CoV-2 variant B.1.1.7 lineage, also known as clade GR from Global Initiative on Sharing All Influenza Data (GISAID), Nextstrain clade 20B, or Variant Under Investigation in December 2020 (VUI – 202012/01), appears to have an increased transmissability in comparison to other variants. Thus, to contain and study this variant of the SARS-CoV-2 virus, it is necessary to develop a specific molecular test to uniquely identify it. Using a completely automated pipeline involving deep learning techniques, we designed a primer set which is specific to SARS-CoV-2 variant B.1.1.7 with >99% accuracy, starting from 8,923 sequences from GISAID. The resulting primer set is in the region of the synonymous mutation C16176T in the ORF1ab gene, using the canonical sequence of the variant B.1.1.7 as a reference. Further *in-silico* testing shows that the primer set’s sequences do not appear in different viruses, using 20,571 virus samples from the National Center for Biotechnology Information (NCBI), nor in other coronaviruses, using 487 samples from National Genomics Data Center (NGDC). In conclusion, the presented primer set can be exploited as part of a multiplexed approach in the initial diagnosis of Covid-19 patients, or used as a second step of diagnosis in cases already positive to Covid-19, to identify individuals carrying the B.1.1.7 variant.

## Introduction

As the pandemic of SARS-CoV-2 continues to affects the planet, researchers and public health teams around the world continue to monitor the virus for acquired mutations that may lead to higher threat for developing COVID-19. Although SARS-CoV-2 mutates with an average evolutionary rate of 10-4 nucleotide substitutions per site each year^1^, a new variant has been recently reported in the UK as a Variant Under Investigation (VUI - 202012/01)^2, 3^, which belongs to the B.1.1.7 lineage^2, 4^, Nextstrain clade 20B^5^, or clade GR from GISAID (Global Initiative on Sharing All Influenza Data)^6^. This variant presents 14 non-synonymous mutations, 6 synonymous mutations and 3 deletions. The multiple mutations present in the viral RNA encoding for the spike protein (S) are of most concern, such as the deletion Δ69-70, deletion Δ144, N501Y, A570D, D614G, P681H, T716I, S982A, D1118H^3, 4^. The SARS-CoV-2 S protein mutation N501Y alters the protein interactions involved in receptor binding domain. The N501Y mutation has been shown to enhance affinity with the host cells ACE2 receptor^4, 7^ and to be more infectious in mice^8^).

Even if the clinical outcomes and additive effects of the mutations present on the B.1.1.7 SARS-CoV-2 variant are still unknown, its rate of transmission has been estimated to be 56%-70% higher, and its reproductive number (Rt) seems to be up to 0.4 higher^2, 9^. The presence of the B.1.1.7 variant has been rapidly increasing in the UK^4^. This and other N501Y carrying SARS-CoV-2 variants have also been identified in other parts of Europe, Australia, USA, Brazil, South Africa, and Egypt^5, 6, 10, 11^.

Several diagnostic kits have been proposed and developed to diagnose SARS-CoV2 infections. Most kits rely on the amplification of one or several genes of SARS-Cov-2 by real-time reverse transcriptase-polymerase chain reaction (RT-PCR)^12, 13^. Recently, Public Health England was able to identify the increase of the B.1.1.7 SARS-CoV-2 variant through the increase in S-gene target failure (negative results) from the otherwise positive target genes (N, ORF1ab) in their three target gene assay^2^. However, to the best of our knowledge, currently no specific test exists or has been developed to identify the B.1.1.7 SARS-CoV-2 variant.

In a previous work^14^, we developed a methodology based on deep learning, able to generate a primer set specific to SARS-CoV-2 in an almost fully automated way. When compared to other primers sets suggested by GISAID, our approach proved to deliver competitive accuracy and specificity. Our results, both *in-silico* and with patients, yielded 100% specificity, and sensitivity similar to widely-used diagnostic qPCR methods. One of the main advantages of the proposed methodology was its ease of adaptation to different viruses, given a sufficiently large number of sequences. In this work we improved the existing semi-automated methodology, making the pipeline completely automated, and created a primer set for the SARS-CoV-2 variant B.1.1.7 in an extremely reduced amount of time (16 hours). The developed primer set, tested *in-silico*, proves to be extremely effective. With this new result, we believe that our method represents a rapid and effective diagnostic tool, able to support medical experts both during the current pandemic, as new variants of SARS-CoV-2 may emerge, and possibly during future ones, as still unknown virus strains might surface.

## Results

As explained in the Methods section, the first step is to run a Convolution Neural Network (CNN) classifier on the data. This yields an average classification accuracy of 99.66% on the testing subset. Secondly, from an analysis of the features constructed by the CNN, we extract 7,127 features, corresponding to RNA subsequences. Next, we ran a state-of-the-art stochastic feature selection algorithm 10 times, to uncover the most meaningful subsequences for the identification of variant B.1.1.7: as shown in Fig. 1, while the best result corresponds to a set of 16 features, using only one is enough to obtain over 99% accuracy, a satisfying outcome for our objective.

**Figure 1.**
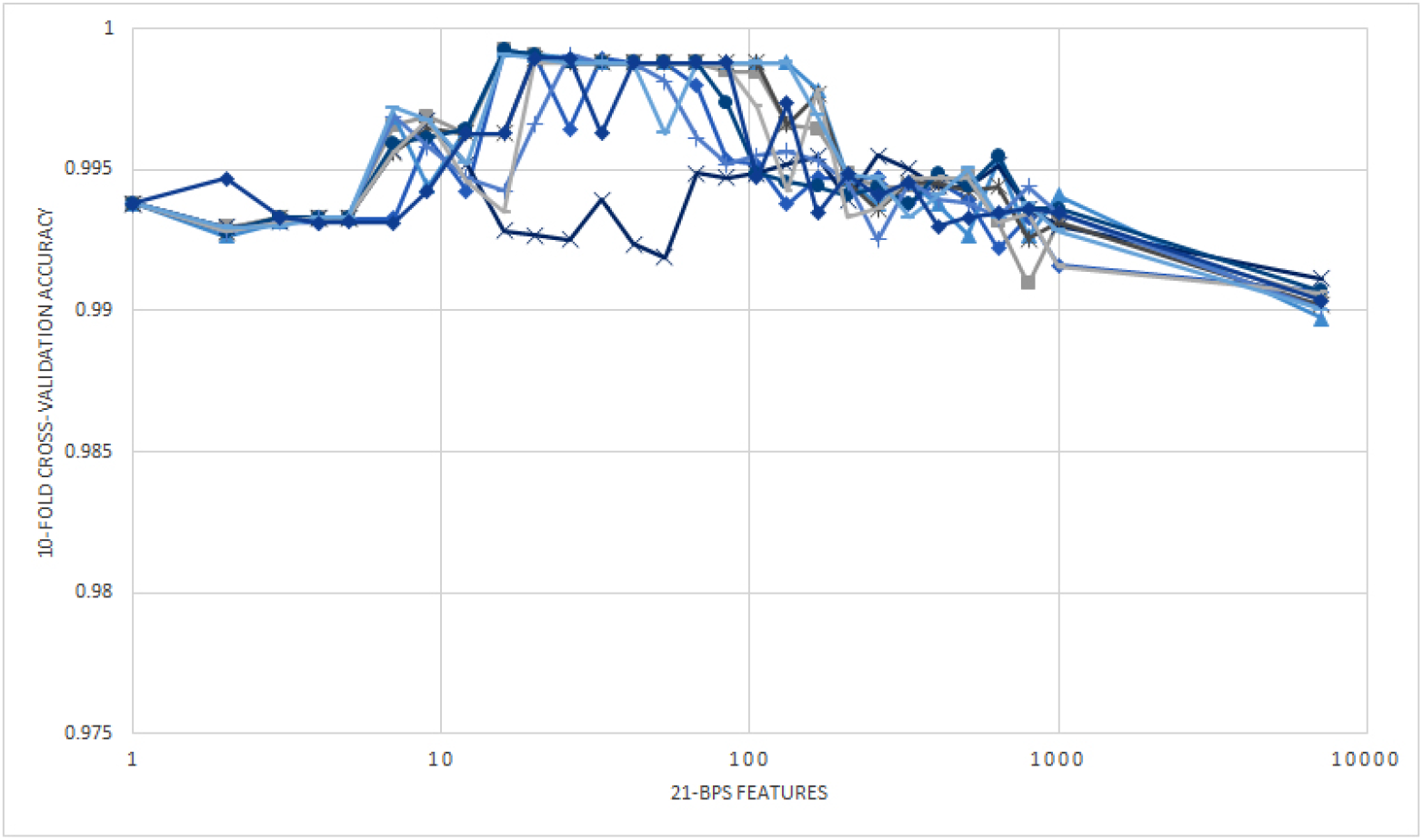
10 runs of the recursive ensemble feature selection algorithm in 803 sequences of training set.

These features are good candidates for forward primers. From the 10 runs, we get 9 different 21-bps features (1 was repeated, found in two different runs): 5 out of the 10 point to mutation Q27stop (C27972T), 3 point to mutation I2230T (T6954C) and 2 to a synonymous mutation (C16176T). Using Primer3Plus we calculate a primer set for each of the 10 features, using sequence EPI_ISL_601443 (see Table 1). From these, only the two features that include mutation C16176T are suitable for a forward primer. The two features are **ACC TCA AGG TAT TGG GAA CCT** and **CAC CTC AAG GTA TTG GGA ACC**: it is easy to notice that the two features are actually part of the same sequence, just displaced by a bps, and therefore generate the same reverse primer **CAT CAC AAC CTG GAG CAT TG** with Tm 58.8°C and 60.6°C respectively for the forward primer, and 60.1°C for the reverse primer.

**Table 1.**
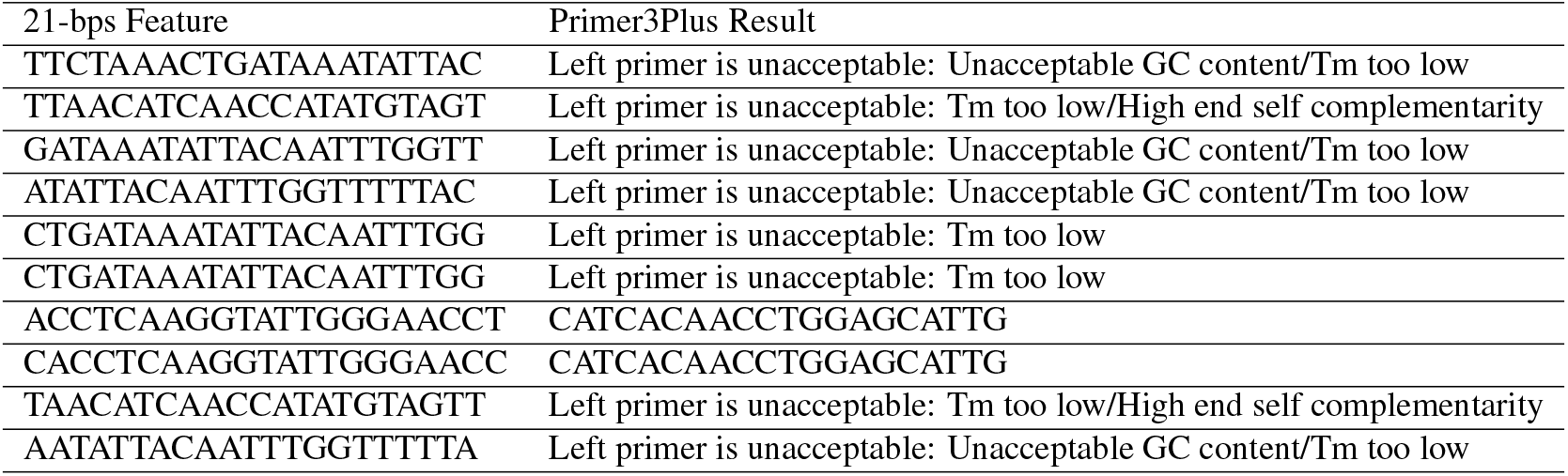
Results from Primer3Plus in sequence EPI_ISL_601443, attempting to use the ten features identified by the feature selection process as primers. Only two features are acceptable primers, and both point to the same sequence.

Using just the feature **ACC TCA AGG TAT TGG GAA CCT**, it is possible to build a simple rule-based classifier that assigns a sample to variant B.1.1.7 if the feature is present. For further validation, we test the rule-based classifier on the 893 sequences of the test set, yielding an area under the curve (AUC) of 0.98, a result considered as an excellent diagnostic accuracy (AUC 0.9-1.0)^15, 16^, see Fig. 2.

**Figure 2.**
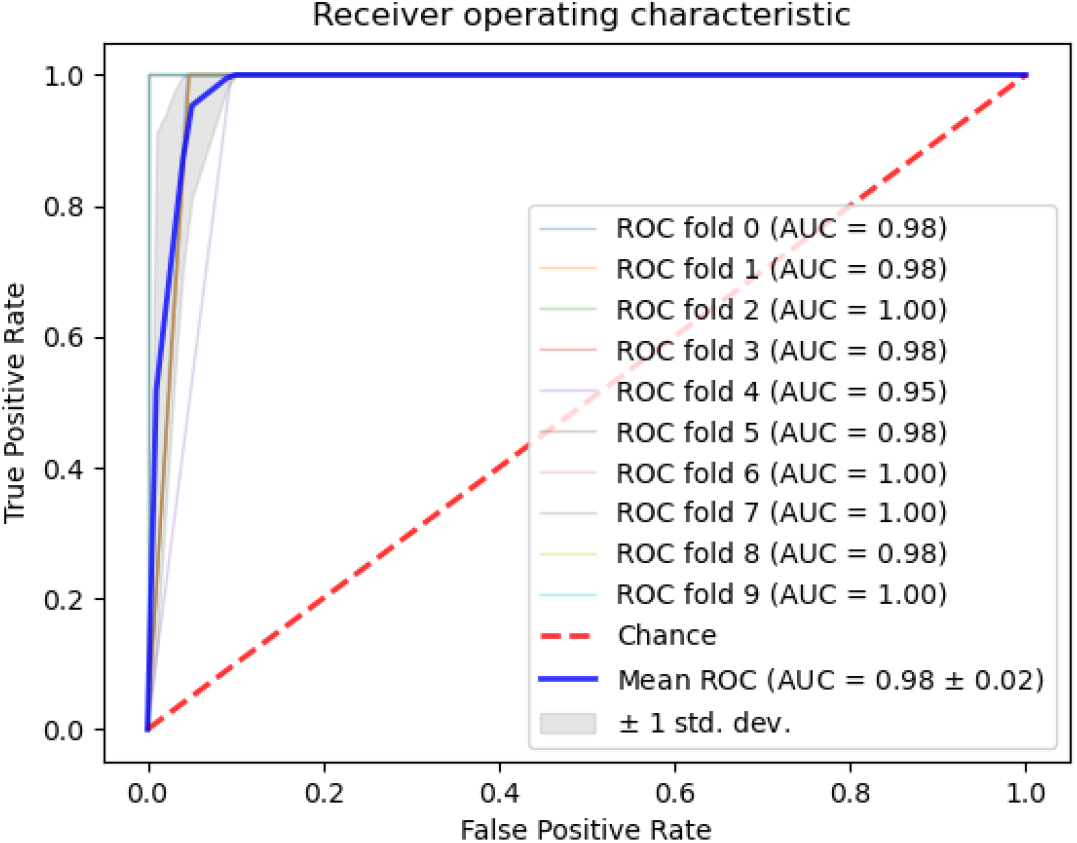
ROC curve of a simple rule-based classifier checking the presence of feature **ACC TCA AGG TAT TGG GAA CCT**, in a 10-fold cross-validation.

We then validate our results on 487 samples of other coronaviruses from the National Genomics Data Center (NGDC)^17^. Checking for the presence of feature **ACC TCA AGG TAT TGG GAA CCT** finds that it is exclusive to B.1.1.7 SARS-CoV-2, with no appearance in any other coronavirus sample. Further validation on 20,571 samples belonging to other viruses from the National Center for Biotechnology Information (NCBI)^18^, shows no appearance of the sequence in any other virus.

## Discussion

The 8,923 SARS-CoV-2 samples downloaded from the GISAID repository show 278 variants besides B.1.1.7. We check the presence of mutations N501Y, A570D, D614G, P681H, T716I, S982A, D1118H, and the forward primer **ACC TCA AGG TAT TGG GAA CCT** among all samples. To verify the presence of mutations, we generated 21-bps sequences, with 10 bps before and after the mutation: for example, mutation N501Y (A23063T) will be **CCAACCCACT T ATGGTGTTGG**. The frequency of appearance of each mutation and the forward primer is reported in Table 2.

**Table 2.** Frequency of appearance of the most significant mutations and the forward primer in the 8,293 sequences from the GISAID dataset.

|  | **B.1.1.7**<br>**# samples (%)** | **Other Variants**<br>**# samples (%)** |
| --- | --- | --- |
| N501Y | 1,985 (94.34%) | 14 (0.02%) |
| A570D | 2,013 (95.67%) | 1 (< 0.01%) |
| D614G | 2,096 (99.62%) | 5,384 (78.96%) |
| P681H | 2,014 (95.72%) | 1 (< 0.01%) |
| T716I | 2,005 (95.29%) | 1 (< 0.01%) |
| S982A | 2,008 (95.43%) | 0 (0%) |
| D1118H | 2,011 (95.57%) | 0 (0%) |
| Forward Primer | 2,007 (95.39%) | 0 (0%) |
| Total Samples | 2,104 (100%) | 6,819 (100%) |

The generated forward primer appears in 2,007 of 2,104 B.1.1.7 sequences, with a frequency of 95.4%. Nevertheless, a further analysis shows that only 2,014 of the sequences labeled as B.1.1.7 present 5 or more of the 7 studied mutations, which can point to an error in annotating the variant in the GISAID dataset, or several extra mutations in the generated 21-bps sequences. If we consider as proper B.1.1.7 variants only sequences that show 5 or more of the mutations, then our primer correctly identifies 2,095 out of 2,104 samples, for a 99.6% accuracy.

Considering we generated 21-bps sequences for the mutations, we can also test them as forward primers using Primer3Plus, which yields **TGA TAT CCT TGC ACG TCT TGA** in spike gene (S982A) as a possible forward primer (see Table 3). The Tm of the forward primer is 59.3°C and the reverse primer sequence will be **GAG GTG CTG ACT GAG GGA AG**.

**Table 3.**
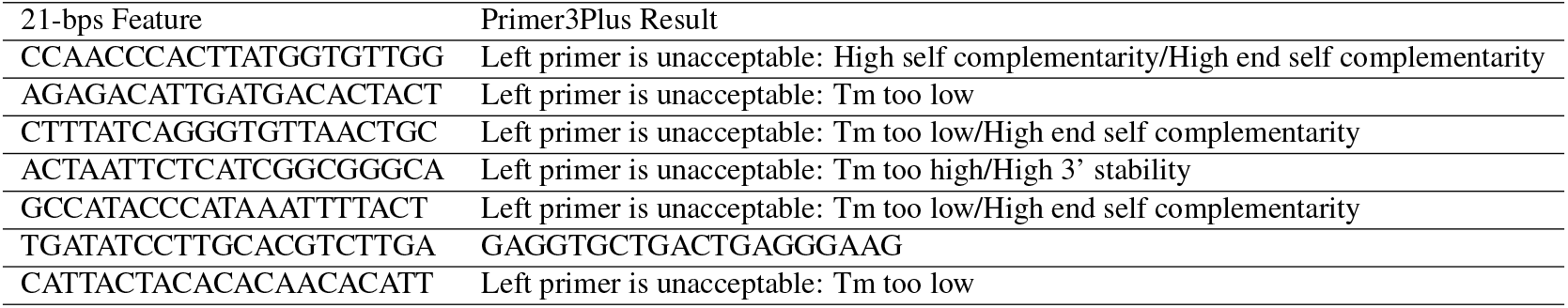
Result of the test of the sequences used to verify the presence of mutations when used as forward primers in sequence EPI_ISL_601443.

A wide variety of diagnostic tests have been used by high-throughput national testing systems around the world, to monitor the SARS-CoV-2 infection^12^. The arising prevalence of new SARS-CoV-2 variants such as B.1.1.7 has become of great concern, as most RT-PCR test to date will not be able to distinguish these new variants because they where not designed for such a purpose. Therefore, public health officials most rely on their current testing systems and their sequencing results to draw conclusions on the prevalence of new variants in their territories. An example of such case has been seen in UK, where they were able to identify the increase of the B.1.1.7 SARS-CoV-2 variant infection in their population only through an increase in the S-gene target failure in their three target gene assay (N+, ORF1ab+, S−) coupled with sequencing of the virus and RT-PCR amplicons products ^2^. Researchers believe that the S-gene target failure occurs due to the failure of one of the RT-PCR probes to bind as a result of the Δ69-70 deletion in the SARS-CoV-2 spike protein, present on B.1.1.7 ^2^. This Δ69-70 deletion, which affects its N-terminal domain, has been recurrently occurring in different SARS-CoV-2 variants around the world^3, 5, 6^ and has been associated with other spike protein receptor binding domain changes^4^. Due to the likeliness of mutation in the S-gene, assays relying solely on its detection are not recommended, and a multiplex approach is required^12, 13, 19^. This is consistent with other existing designs like CoV2R-3 in the S-gene^20^, that will also yield negative results for the B.1.1.7 variant, as the reverse primer sequence is in the region of mutation P681H. A more in-depth analyisis of S-dropout positive results can be found in Kidd et al. ^21^.

Given the concern of the increase in prevalence of the new variant SARS-CoV2 B.1.1.7 and its possible clinical implication in the ongoing pandemic, diagnosing and monitoring the prevalence of such variant in the general population will be of critical importance to help fight the pandemic and develop new policies. In this work, we propose 2 possible primer sets that can be used to specifically identify B.1.1.7 SARS-CoV2 variant. We believe that our primers can be used in a multiplexed approach in the initial diagnosis of Covid-19 patients, or used as a second step of diagnosis in cases already verified positive to SARS-CoV-2 to identify individuals carrying the B.1.1.7 variant. In this way, health authorities could then better evaluate the medical outcome of this patients, and adapt or inform new policies that can help curve the rise of variants of interest. Although the proposed primer sets delivered by our automated methodology will still require laboratory testing to be validated, our deep learning design can enable the timely, rapid, and low-cost operations needed for the design of new primer sets to accurately diagnose new emerging SARS-CoV-2 variants and other infectious diseases.

## Methods

From the GISAID repository we downloaded 10,712 SARS-CoV-2 sequences on December 23rd, 2020. After removing repeated sequences, we obtain a total of 2,104 sequences labeled as B.1.1.7, and 6,819 sequences from other variants, for a total of 8,923 samples. B.1.1.7 variants are assigned class label 0, while all the remaining samples are labeled as class 1.

Following the procedure described in Lopez et al.^14^, there are 4 steps for the automated design of a specific primer for a virus: (i) run a CNN for the classification of the target virus against other strains, (ii) translate the CNN weights into 21-bps features, (iii) perform feature selection to identify the most promising features, and (iv) carry out a primer simulation with Primer3Plus^22^ for the features uncovered in the previous step. While in^14^ the proposed pipeline was only partially automatic, and still required human interventions between steps, in this work all steps have been automatized, and the whole pipeline has been run with no human interaction. The experiments, from downloading the sequences to the final *in-silico* testing of the primers, took around 16 hours of computational time on a standard end-user laptop.

In a first step, we train a convolution neural network (CNN) using 8,030 sequences for training and 893 for testing. The architecture of the network is shown in Fig 3, and is the same as the one previously reported in^14^. Next, as the classification accuracy of the CNN is satisfying (>99%), using a subset of 803 training sequences we translate the CNN weights into 21-bps features, necessary to differentiate between B.1.1.7 variant samples and the others. The length of the features is set as 21 bps, as a normal length for primers to be used in RT-PCR tests is usually 18-22 bps. Then, we apply recursive ensemble feature selection^23, 24^ to obtain the most meaningful features that separate the two classes. Finally, we simulate the results of using the most promising features obtained in the previous step as primers, using Primer3Plus^22^ in the sequence EPI_ISL_601443, that is the canonical variant of concern^2^.

**Figure 3.**
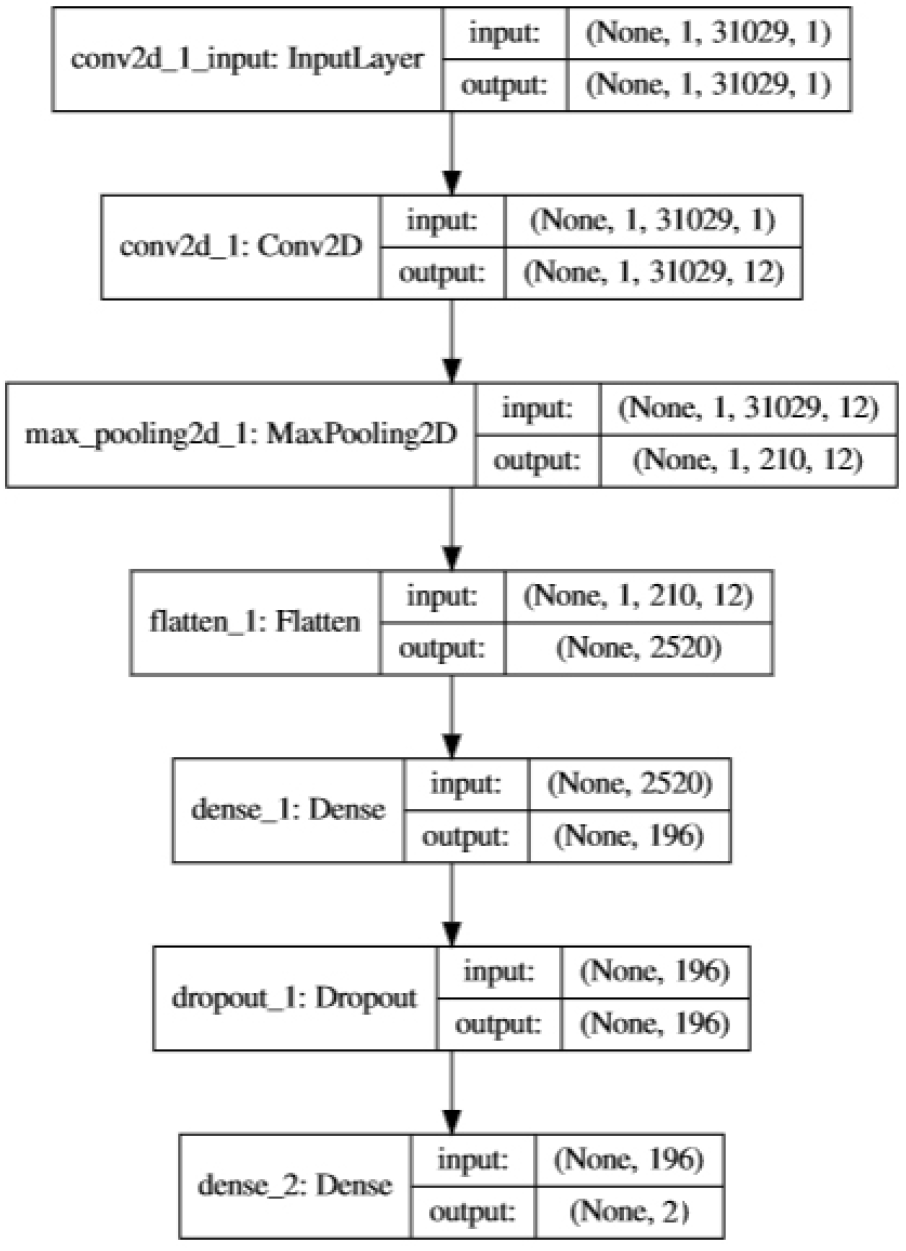
CNN Architecture to identify Variant B.1.1.7.

For further *in-silico* analysis, the resulting features will be compared against 487 sequences of other coronaviruses from the NGDC repository, to check if our generated primers are specific enough to SARS-CoV-2 (Table 4). In addition, we used data from NCBI with 20,571 viral samples from other taxa, for a total of more than 584 other viruses (not considering strains and isolates). The whole procedure is summarized in Fig. 4.

**Table 4.**
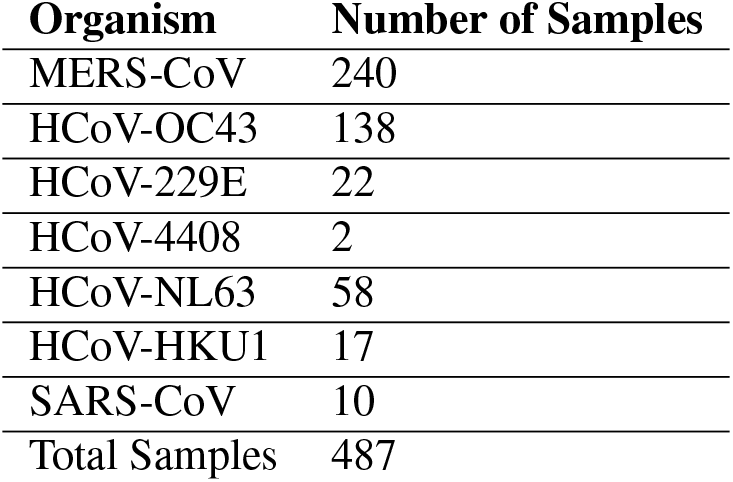
Organism and number of samples of other coronaviruses to compare specificity in the sequences.

**Figure 4.**
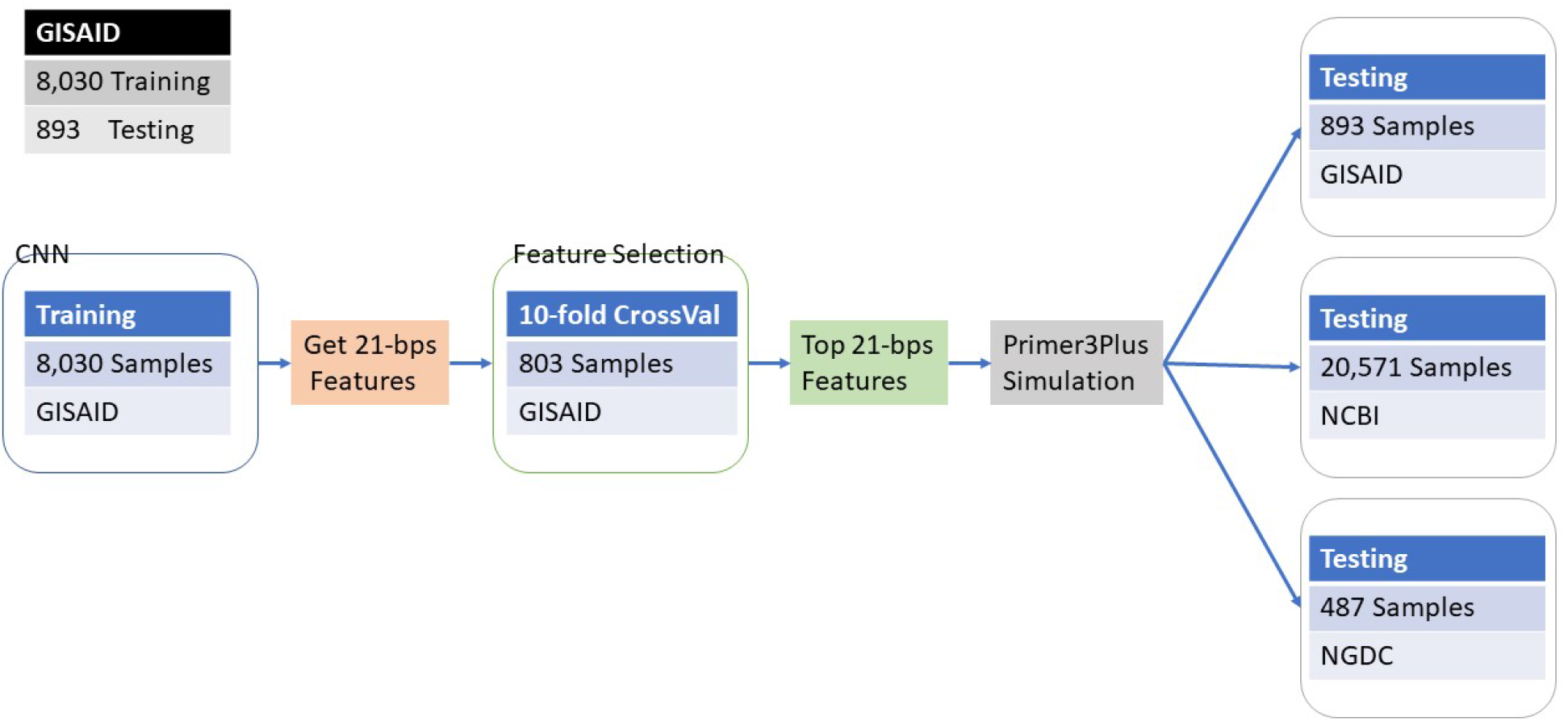
Summary of construction of the primers and validation.

## Additional information

The authors declare no competing interests.

## Author contributions statement

CAP, LMM, made the biological analysis, and primer design. ALR and AT made the programming, data collection and experiments in silico. EC, ADK and JG made the experiment and study design. All the authors contributed to the writing.

